# Temporal landscape of mutation accumulation in SARS-CoV-2 genomes from Bangladesh: possible implications from the ongoing outbreak in Bangladesh

**DOI:** 10.1101/2020.08.20.259721

**Authors:** Otun Saha, Rokaiya Nurani Shatadru, Nadira Naznin Rakhi, Israt Islam, Md. Shahadat Hossain, Md. Mizanur Rahaman

## Abstract

Along with intrinsic evolution, adaptation to selective pressure in new environments might have resulted in the circulatory SARS-CoV-2 strains in response to the geoenvironmental conditions of a country and the demographic profile of its population. Thus the analysis of genomic mutations of these circulatory strains may give an insight into the molecular basis of SARS-CoV-2 pathogenesis and evolution favoring the development of effective treatment and containment strategies. With this target, the current study traced the evolutionary route and mutational frequency of 198 Bangladesh originated SARS-CoV-2 genomic sequences available in the GISAID platform over a period of 13 weeks as of 14 July 2020. The analyses were performed using MEGA 7, Swiss Model Repository, Virus Pathogen Resource and Jalview visualization. Our analysis identified that majority of the circulating strains in the country belong to B and/or L type among cluster A to Z and strikingly differ from both the reference genome and the first sequenced genome from Bangladesh. Mutations in Nonspecific protein 2 (NSP2), NSP3, RNA dependent RNA polymerase (RdRp), Helicase, Spike, ORF3a, and Nucleocapsid (N) protein were common in the circulating strains with varying degrees and the most unique mutations(UM) were found in NSP3 (UM-18). But no or limited changes were observed in NSP9, NSP11, E (Envelope), NSP7a, ORF 6, and ORF 7b suggesting the possible conserved functions of those proteins in SARS-CoV-2 propagation. However, along with D614G mutation, more than 20 different mutations in the Spike protein were detected basically in the S2 domain. Besides, mutations in SR-rich region of N protein and P323L in RDRP were also present. However, the mutation accumulation showed an association with sex and age of the COVID-19 positive cases. So, identification of these mutational accumulation patterns may greatly facilitate drug/ vaccine development deciphering the age and the sex dependent differential susceptibility to COVID-19.

## 1. Introduction

In the past two decades, Coronaviruses mainly of the β-coronavirus family *Coronaviridae* and the subfamily *Coronavirinae* have been a major subject of deeper investigations due to their emergence, re-emergence and associated public health impact (Mim et al., 2020; Saha et al., 2020). Among the seven coronaviruses (229E, OC43, NL63, HKU1, SARS-CoV (Severe Acute Respiratory Syndrome Coronavirus), MERS-CoV (Middle East respiratory syndrome Coronavirus) and SARS-CoV-2 responsible for coronavirus disease 2019 (COVID-19)) causing human infections, the newly emerged single-stranded RNA beta-coronavirus SARS-CoV-2 has been wreaking havoc around the world since its emergence in mid-December 2019 in the Chinese city of Wuhan (Rahaman et al.,2020; Mim et al., 2020; Saha et al., 2020) and was first reported from Bangladesh on March 8, 2020.

This 29903 kb enveloped virus consists of a 5′□untranslated region (5′□UTR), spike (S), envelope (E), matrix (M), nucleocapsid (N) gene and 3′□UTR (Fani et al., 2020), among which E, M, and N proteins function to protect the viral genome and S protein binds to the host cell receptor (Kaushal et al., 2020). On the other hand, among the sixteen non-structural proteins (NSPs), RNA dependent RNA polymerase (RdRp) (NSP12), helicase (NSP13), mRNA capping (NSP14 and NSP16), and fidelity control (NSP14) synthesize and process the viral RNA (Kaushal et al., 2020), while the remaining proteins are crucial cofactors facilitating the function of viral enzymes (da Silva et al., 2020). So, the current circulating strain might have evolved through the ongoing evolutionary process of mutations in these genes since its emergence (Dawood, 2020). Errors made by RdRp despite having proofreading activity (Sevajol et al., 2014) along with a direct response to selective pressure on the viral genome and homologous recombination may lead to mutational accumulation in the SARS-CoV-2 genome (Hon et al., 2008), while according to recent studies on mutation analysis, no recombination events were reported (Yu, 2020) and the sequence diversity of SARS-CoV-2 so far is very low (Fauver et al., 2020). On the contrary, the receptor-binding domain (RBD) in the S protein is the most variable genomic part in the betacoronavirus group (Wu et al., 2020; Zhou et al., 2020), and some sites of S protein might be subjected to positive selection (Lv et al., 2020). However, despite these variabilities in the SARS-CoV-2 genome, one key question remains as to whether these mutations have any functional impact on the pathogenicity of SARS-CoV-2. The previous experiences with MERS-CoV and SARS-CoV (Tang et al., 2014) the close relatives of SARS-CoV-2 (Wu et al., 2020; Zhou et al., 2020), showed that single mutation might be significant enough to confer resistance to neutralizing antibodies against those viruses. Meanwhile, during the rampant spread of SARS-CoV-2 around the world, it has undergone multiple antigenic drifts including several mutations compromising the containment and diagnostics strategies along with the effectivity of repurposed drugs (Coppee et al., 2020), which suggested that the virus will be active and spreading for a year or more before vaccines are available (Wu et al., 2020; Zhou et al., 2020). Besides, based on amino acid changes of the genomes, 3 major clades (S, G, and V) were proposed in many more studies (Wu et al., 2020; Zhou et al., 2020; Ceraolo et al., 2020). Another study by Tai et al. in 2020 suggested that amino acid variations in the genome are associated with the stability of RBD/ACE2 structure. Also, the primer-template mismatches might affect the stability and the functionality of polymerase (Kim, et al., 2020). On the other hand, a study by Su et al. in 2020 revealed that the deletion of 382 nucleotides towards the 3’ end of the viral genome may have an impact on the viral phenotype. Thus, these mutational analyses justify the potential of mutations in affecting the viral infectivity and adaptability to the new environment as well as explaining the differential rates of infection and mortality worldwide conducive to controlling the pandemic. Meanwhile, the data avalanche, especially the complete genome sequences in Global Initiative on Sharing All Influenza Data (GISAID, https://www.gisaid.org/) has resulted in an unprecedented expeditious effort towards understanding the implications of genome diversity (Andersen et al., 2020; Shen et al., 2020) in pathogenicity, drug repositioning (Wu et al., 2020; Zhou et al., 2020) or developing diagnostic and preventive strategies (Zhao and Chen, 2020). Concurrently with the global sequence data, legionary complete genome sequences have been submitted from Bangladesh in GISAID since the first submission on 14 July, 2020 (Islam et al., 2020). So, the current study was designed to investigate the genomic diversity of SARS-CoV-2 strains isolated from the country as well as analyzing the temporal profile of the mutational accumulations in the genome. Ultimately, this study will give an insight into the circulating strains of the country to devise a more effective containment strategy and efficient treatment regimen along with adding values to the global understanding of SARS-CoV-2 genome evolution and molecular basis of its pathogenicity, infectivity and drug/vaccine targets.

## 2. Materials and Methods

### 2.1 Retrieval of SARS-CoV-2 genome sequences from the database

SARS-CoV-2 genome analysis reveals the genomic variation of the sequence that causes alterations in protein encoding. A total of 226 complete genome sequences of SARS-CoV-2 isolated from Bangladesh were retrieved from the GISAID virus database (https://www.gisaid.org/, last access 14 July 2020) along with the collection date and the patient history (Supplementary Material SM1 & Supplementary Table ST1). Multiple sequence alignment (MSA) was performed using a virus pathogen resource (https://www.viprbrc.org/) followed by MEGA 7 (Kumar et al., 2016) to remove ambiguous and low-quality sequences. Later, the MSA file was opened with Jalview visualization software to eliminate the redundancy of the studied sequences (Waterhouse et al., 2009). Finally, the complete viral genomes sequenced from both male and female patients, reported from Bangladesh were analyzed using the reference genome (NC_045512.2).

### 2.2 Analysis of the retrieved genome sequence of the SARS-CoV-2

The nucleotide and amino acid position of each protein of the SARS-CoV-2 genome was locatedusingtheSwissModelRepository (https://swissmodel.expasy.org/repository/species/2697049)andtheGISAID (https://www.gisaid.org/). The genome analysis was initially performed using a free web-based tool, the Virus Pathogen Resource (https://www.viprbrc.org/) which performed phylogenetic analysis to identify clusters present in diverse genome sequences of SARS-CoV-2 isolated from Bangladesh. Also, it facilitated the identification of the type of Coronavirus and the genotype of a protein sequence. The FASTA files of nucleotide sequences retrieved from GISAID were given as input in the tool to get mutational information of the questioned genome using the reference sequence of the virus isolated from the Wuhan seafood market (NC_045512). Nucleotide and protein mutation examination were accomplished manually using GISAID. Mutation frequency for nucleotide and amino acid changes were calculated for each week. The ratio of the total number of mutations in each week and the total number of genomes obtained in that week was used to calculate the mutation frequency.

### 2.3 Phylogenetic Analysis of SARS-CoV-2 sequences

To infer the evolutionary relationships among the examined sequences, the sequences were aligned with relevant reference sequences retrieved from NCBI database using the neighbor-joining approach (Rahman et al., 2020). The Molecular Evolutionary Genetics Analysis across Computing Platforms (MEGA 7) (Kumar et al., 2016) software was used to construct phylogenetic tree applying the neighbor-joining method (Saha et al., 2020) and evolutionary distances were computed using the Kimura-Nei method (Saitou, 1987). The percentage of replicate trees in which the associated taxa clustered together in the bootstrap test (1000 replicates) is shown next to the branches.

## 3. Results

### 3.1 Genome Analysis

A total of 226 complete SARS-CoV-2 genome sequences isolated from the Bangladeshi patients submitted between 18 April and 14 July 2020 and in which majority were B and/or L type (SM1 & ST1) were included in the analysis. Using Jal view visualization, redundant sequences were identified and removed from the analysis. Genome sequences with legionary characters other than A, T, G, and C such as N, R, X, and Y, representing sequencing errors, such as unspecified or unknown nucleotide, unspecified purine nucleotide, and unspecified pyrimidine nucleotide, respectively were also excluded from the analysis. Additionally, sequences with unknown dates of collection or without complete patient’s history were also removed. After considering all the exclusion criteria, 198 unique SARS-CoV-2 genome sequences remained to use for mutational analysis (ST1 & SF1). The complete sequence of all 198 genomes sequences is provided in a SM1, while the accession numbers along with collection dates, patient’s history and mutational status of these sequences are summarized in ST1. The phylogenetic tree of 198 complete SARS-CoV-2 genomic sequences from Bangladesh and the reference sequence (NC_045512) of the isolate from Wuhan, China shows the SARS-CoV-2 sequences from Bangladesh differ from the reference sequence, NC_045512 (mark as blue) of Wuhan, China. Also, the SARS-CoV-2 Bangladesh lineage is split into many sub-lineages, which are represented by different branches. Each branch of the tree ends with a cluster or a single sequence. Closely related genome sequences having a minimum branch deviation (cut of 0.00005) were grouped in clusters (cluster A to Z). As an example, cluster A is formed by six closely related genome sequences from June 2020 (EPI_ISL_483627 (12-06-2020), EPI_ISL_483627 (16-06-2020), EPI_ISL_483635 (17-06-2020), EPI_ISL_4836289 (27-06-2020), EPI_ISL_483689 (26-06-2020), and EPI_ISL_4836214 (14-02-2020). Although some genome sequences were isolated during the different months (April or May or July), they were found closely related and, therefore, grouped in the same cluster (cluster B, W, X, Z). The phylogenetic tree revealed that the genome of more closely related sequences has a low evolutionary rate. Also, the evolution pattern suggests that all lineages share the same ancestry as the Wuhan virus with multiple gene mutations over time. Figure 1 also demonstrates that all sequences of the month of June were more distant from the sequence of Wuhan. Besides, the majority of the sequences were also more distant from the 1st Bangladeshi reported sequence (mark as red) (Figure 1). However, our findings support the notion that the Bangladeshi SARS-CoV-2 virus genome is the product of the Wuhan SARS-CoV-2 evolution. Also, it appears that the virus genome is continuously changing and that could be a result of adaptation to the new environment.

**Figure 1.**
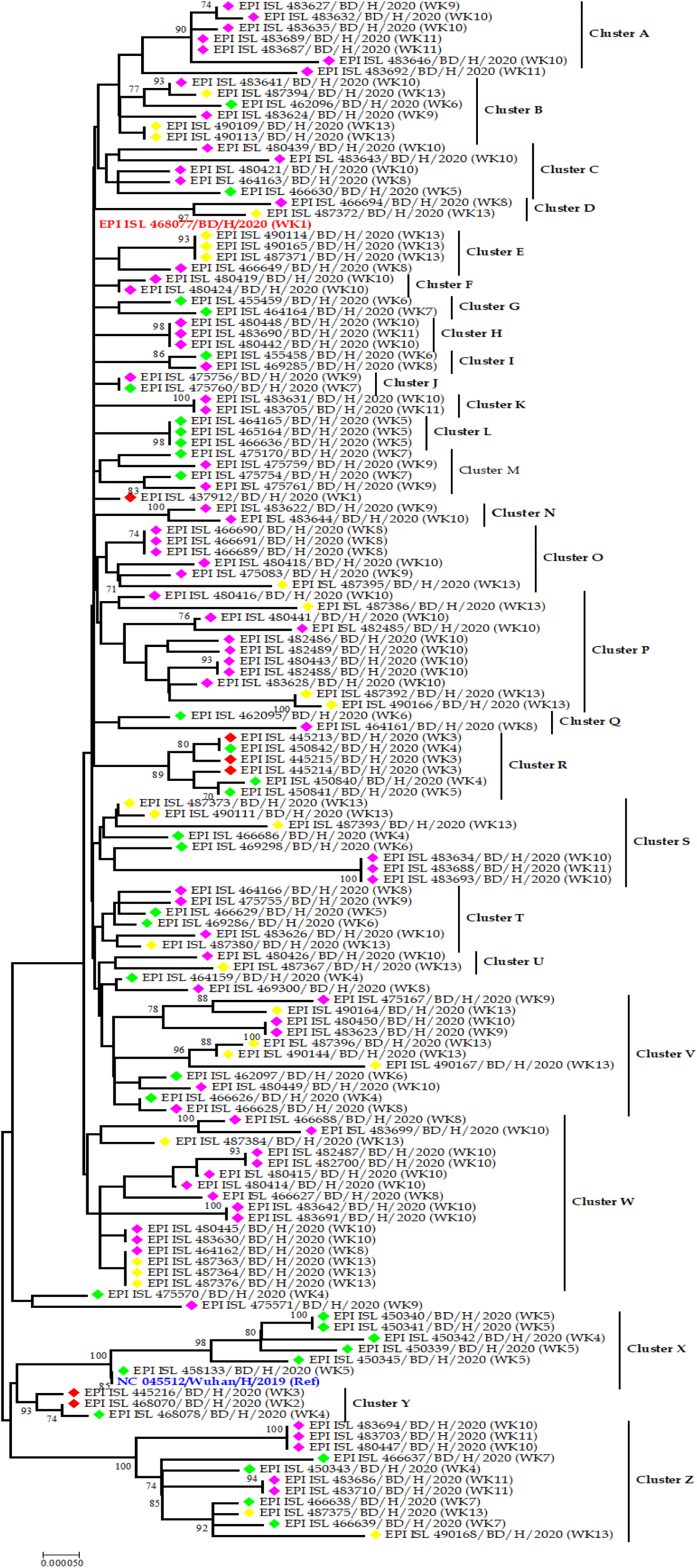
Phylogenetic tree of the studied whole genome sequences of SARS-CoV-2 Bangladesh outbreak. The optimal tree with the sum of branch length = 0.01209279 is shown. The tip of branches corresponds to the accession numbers with country originated, sources, released year and week of sequences. The taxon colored with red, green, pink and yellow for denoting April, May, June and July month respectively. Closely related genome sequences with minimum branch deviation (cut of 0.00005) were represented in clusters (cluster A to Z). Reference sequence (NC_045512) form Wuhan, China and 1^st^ declared sequences form Bangladesh were marked as blue and red respectively. There were a total of 29011 positions in the final dataset. Evolutionary analyses were conducted in MEGA 7. The tree reveals the history of the common ancestry of all 198 SARS-CoV-2 genome sequences from Bangladesh outbreak. The lines of a tree represent evolutionary lineages. Sequences were grouped by the taxon and shown as red, green, pink and yellow mark colors for April, May, June and July, respectively.

### 3.2 Mutational Analysis in SARS-CoV-2 Genome

Mutations were grouped by the date and divided into seven days period, making a total of 13 weeks (Supplementary Table ST2). Because of the presence of one genomic sequence (EPI_ISL_468077) during the 1st week, this sequence was considered with week 2 and declared this 2 weeks as 1st week (W1). To identify the regions with the most mutations over time, SARS-CoV-2 genome was divided into five regions of approximately 6.5 kb each and these were named as R1 (1–6500 bp), R2 (6501–13024 bp), R3 (13025–19620 bp), R4 (19621–26220 bp), and R5 (26221–end) (Figure 2). This was done to facilitate the analysis of the large genome of the SARS-CoV-2 virus, which is 29,903 kb. Protein nonsynonymous mutation frequency was calculated by taking the ratio of the number of total protein mutations and the number of genome sequences in each week. Mutation frequency for whole Genomes was observed to be low during the initial five weeks except region 2. However, after the first five weeks, the frequency seems to have increased sharply until week 9 and remained similar or even slightly down in weeks 10 and 11 (Figure 3). After week 11, all the regions except R4 seems to be increasing sharply again. It appears that the mutational frequency of regions 1, region 4, and region 5 increased over time with a higher frequency in weeks 7–9 (Figure 3). Overall, the mutation frequency during the entire period of analysis (13 weeks) was found to be the highest in region 5, followed by regions 4, and 2 (Figure 3). Regions 2 and 3 appeared to be more conserved. These studied positions of viral proteins were identified using the Swiss model and Genbank. Additionally, the amino acid mutations were attained from the GISAID. The amino acid mutations frequencies were calculated and analyzed over these 13 weeks for all five regions of SARS-CoV-2. The amino acid mutation frequency for each protein is demonstrated in Figure 4. The proteins such as N (Nucleocapsid) and S (Spike) appear to have the highest mutation rates over these 13 weeks (Figure 4). NSP2, NSP3, RdRp (RNA dependent RNA polymerase), ORF3a, also showed substantially high mutation frequency. Interestingly, some proteins such as NSP9, NSP11 had no mutation frequency over the study period. Additionally, proteins E (Envelope), NSP7a, ORF 6, and ORF 7b had the lowest mutational frequency. Unique mutations were also calculated and summarized in Table 1. The total number of unique mutations accumulated during the entire 13 weeks were found to be highest in region 1 followed by region 4 and 5 shown in figure 5. Unique mutation frequency for whole genomes was observed to be highest during the week 9 to 11. However, after the first four weeks, the frequency seems to have increased drastically followed by fluctuating until week 9. After week 11 it’s drastically down in week 12. Most interestingly, after week 12 unique mutation in the region 1 and 5 fell to approximately zero but the mutations in the other three regions increased (Figure 2). Overall, the unique mutation frequency during this entire period of analysis (13 weeks) was found to be highest in region 1 but regions 2 and 3 had appeared to be more conserved as like as normal mutational frequency. Additionally, the total number of the unique mutations identified in different proteins over these 13 weeks study is presented in Table 1. The highest numbers of unique mutations were found in NSP3 (18) followed by S and ORF3a (9) (Table 1). NSP2, RdRp, helicase, E and N proteins had unique mutations ranging between 2 and 7 amino acids per protein (Table 1 and Figure 5). NSP8, NSP9, NSP11, 2’-O-ribose methyltransferase, Matrix, ORF7b had no unique mutations. Certain mutations were identified to be sustained for several weeks, for example, I120F, T412I, L37F, P323L, G204R, R203K, and D614G which were sustained for more than six weeks (table 1 and Figure 5).

**Table 1:**
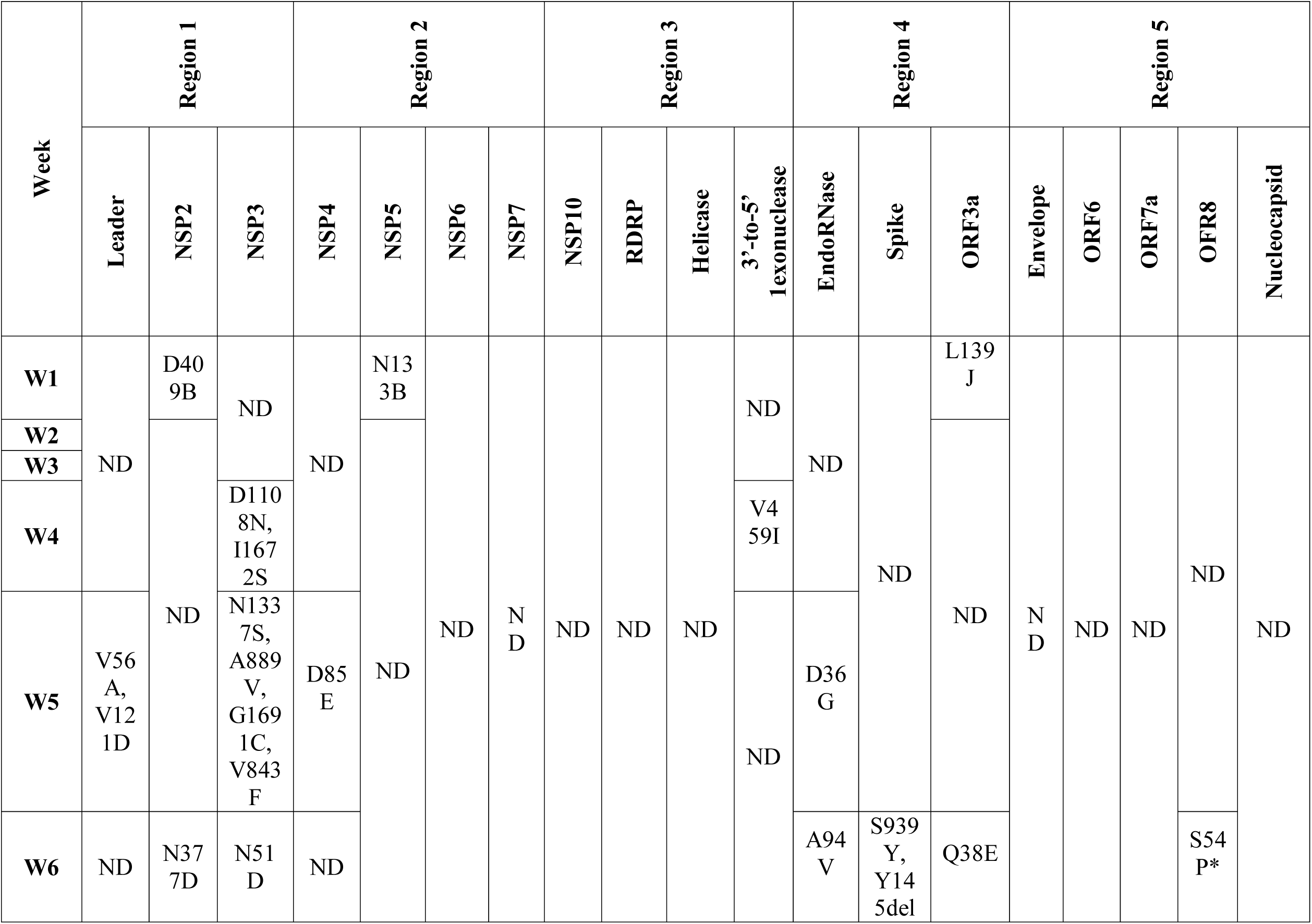
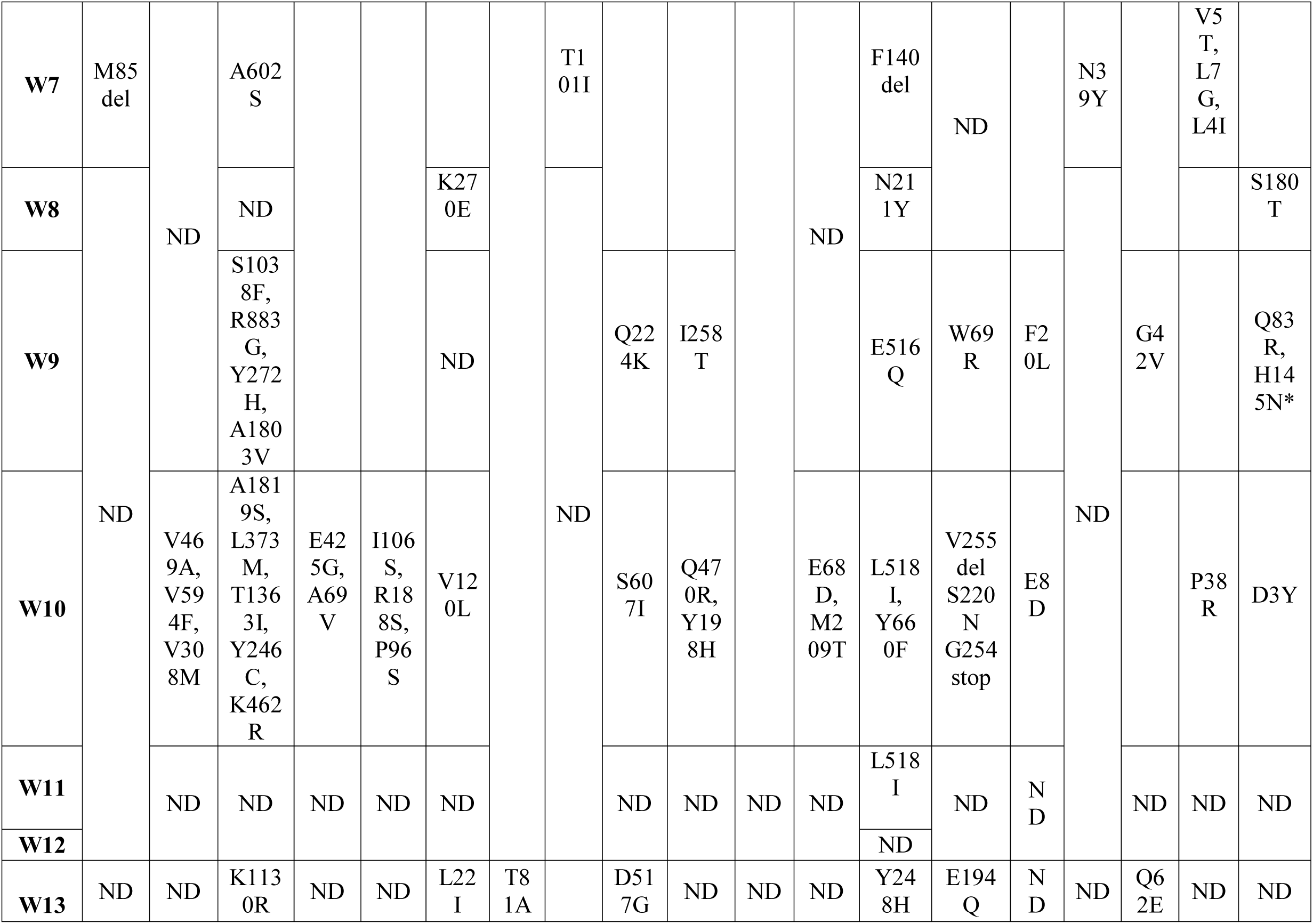
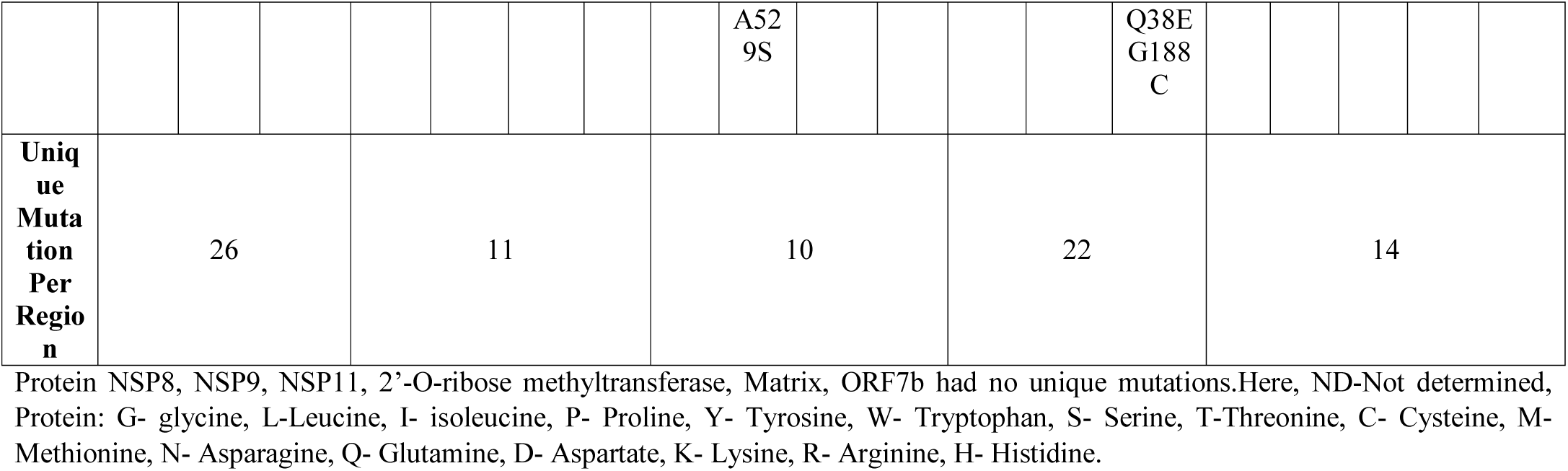
Unique Mutations of five regions occurred in more than one week in SARS-CoV-2 virus. Unique mutations were identified by removing redundant mutations that occurred in more than one week.

**Figure 2.**
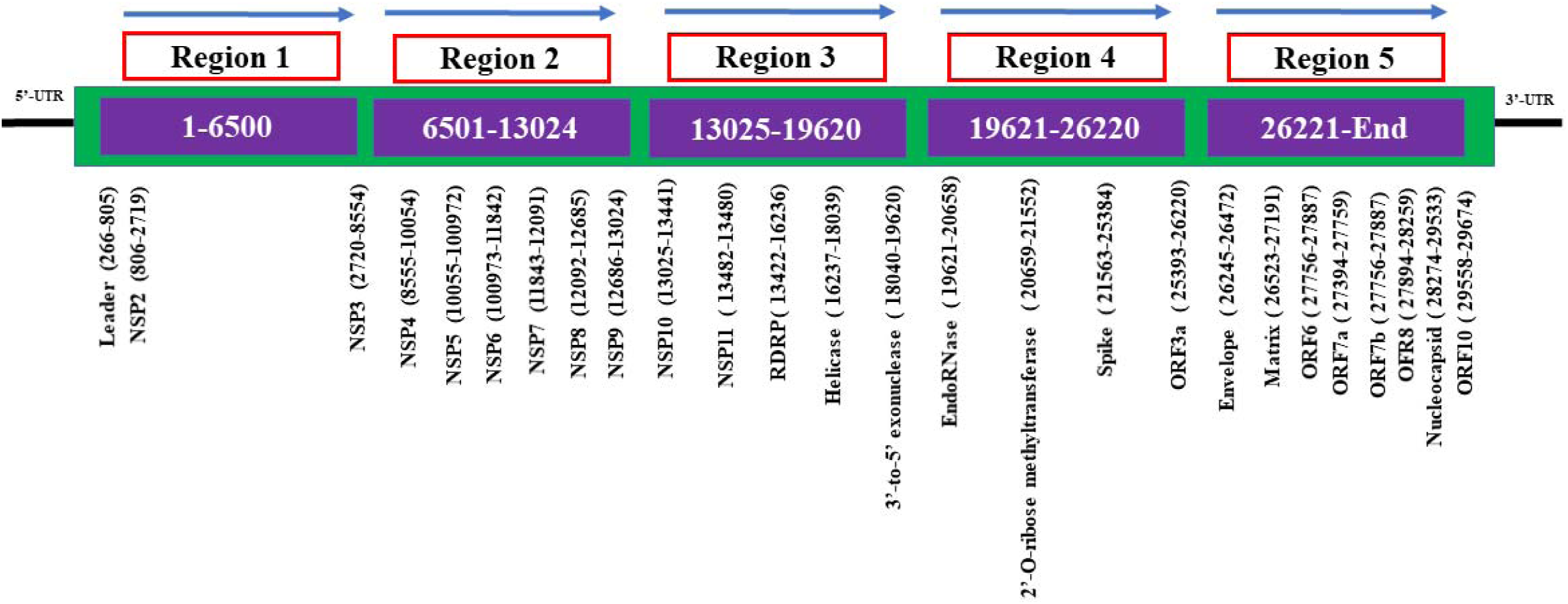
Mapping of SARS-CoV-2 genome regions and proteins. The SARS-CoV-2 genome was divided into five regions and the location of each protein in the different regions i schematically presented.

**Figure 3.**
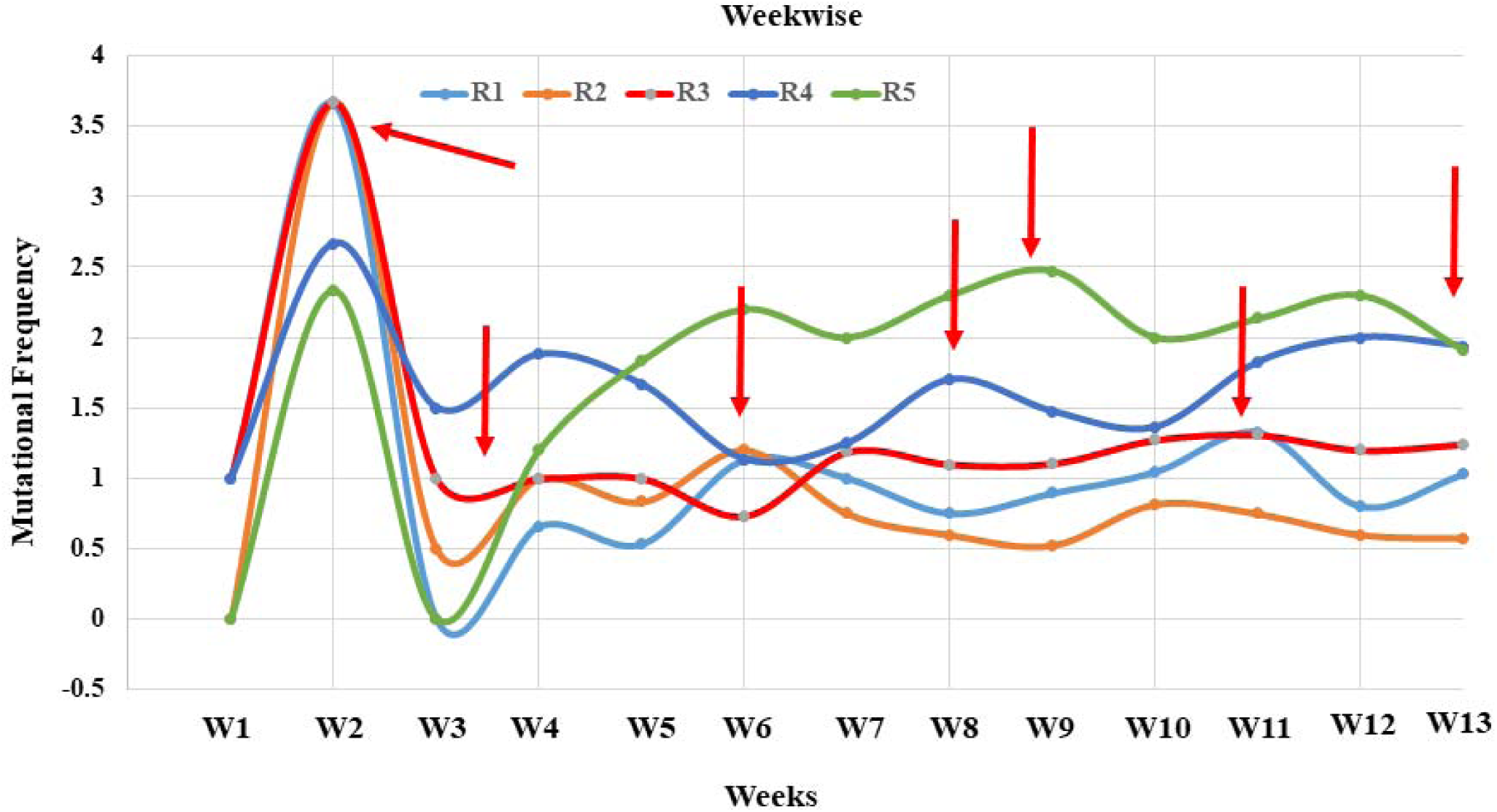
Mutational frequency of five genomic segments of SARS-CoV-2. Mutational frequency was calculated by the ratio of the number of total protein mutations and the number of genome sequences in each week. The SARS-CoV-2 genome was divided into five regions, which are represented as R1–R5.

**Figure 4.**
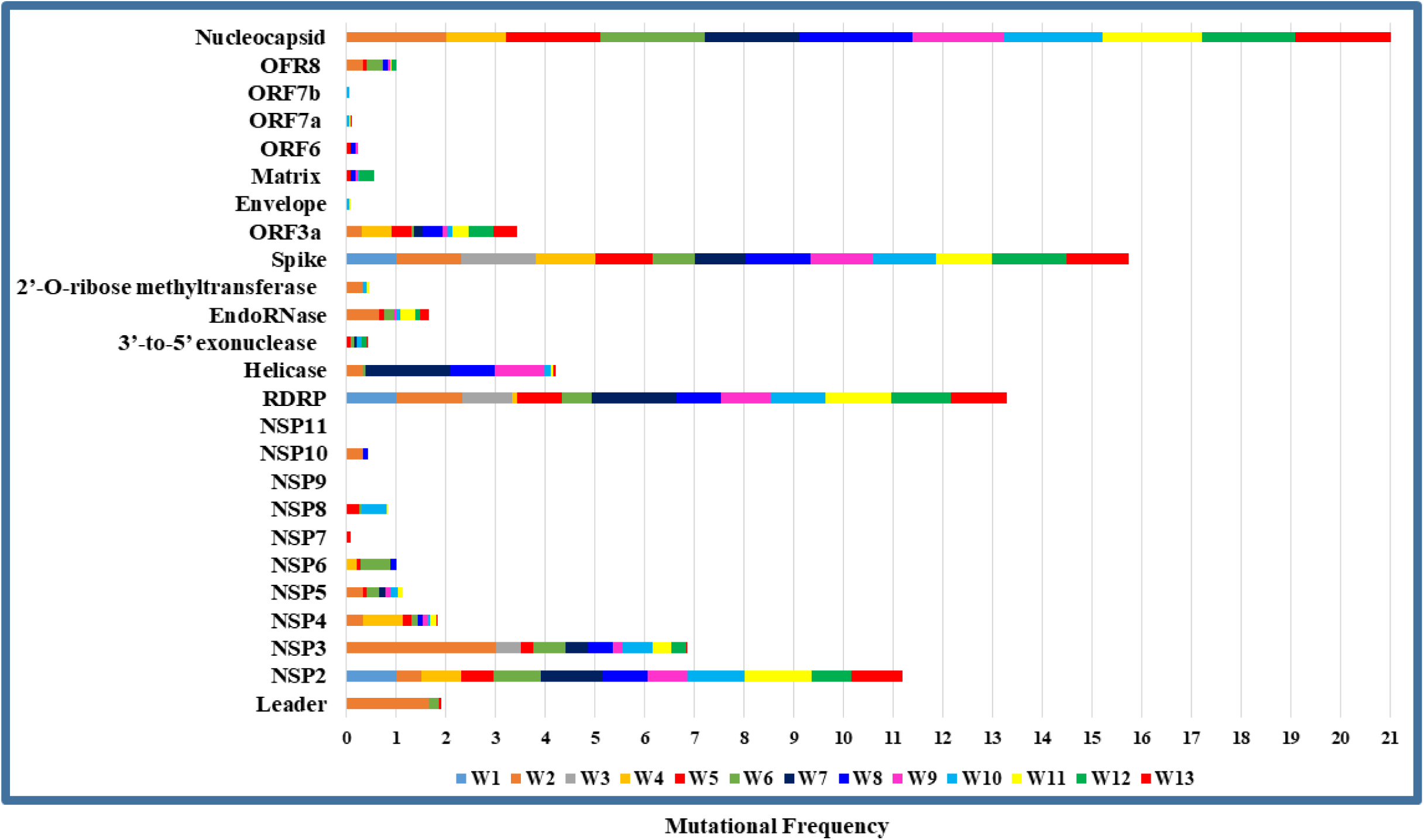
Week wise comparative amino acid mutational frequency of SARS-CoV-2 proteins. Mutational frequency was calculated by the ratio of the number of total amino acid mutations and the number of genome sequences in each week. W1–W13 represents different weeks.

**Figure 5.**
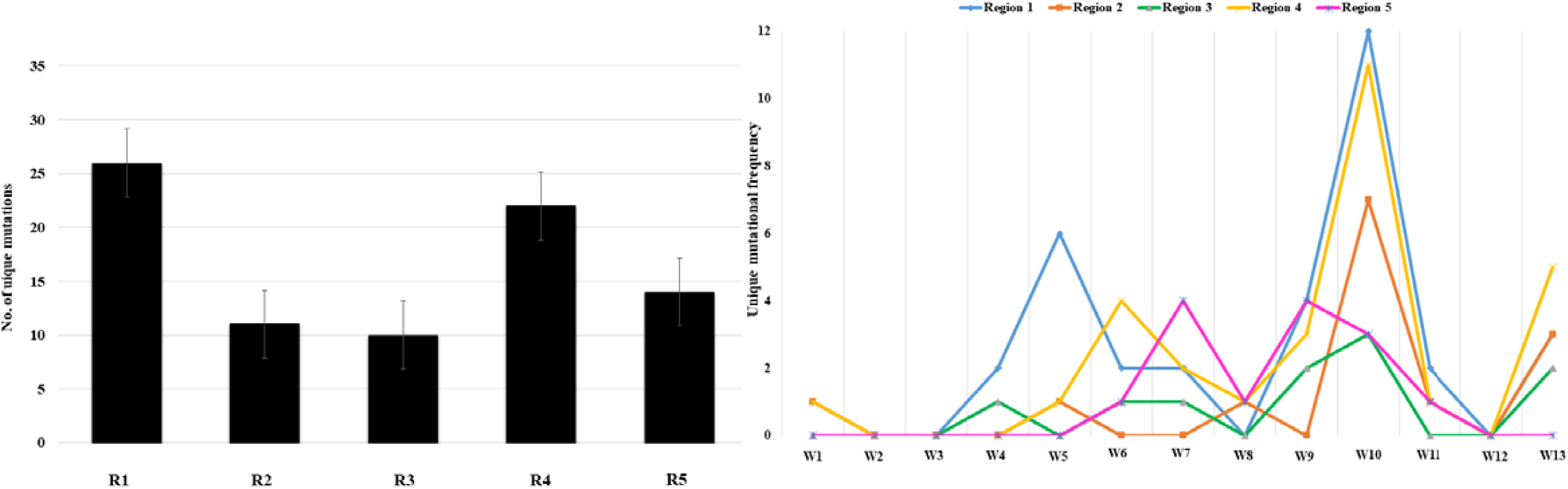
Unique protein mutation of five genomic regions of SARS-CoV-2. A. Region wise unique mutation distributions in the studied genomes. Unique mutations are calculated by removing the redundant mutations, which occur in more than one week. The SARS-CoV-2 genome was divided into five regions, which are represented as R1-R5.B. UNIQUE mutational frequency of five genomic segments of SARS-CoV-2. Mutational frequency was calculated by the ratio of the number of total protein mutations and the number of genome sequences in each week.

### 3.3 Variability of the Spike protein

Spike Protein (S) helps the virus to enter into the host cell, which gets cleaved by the host Proteases at two sites (685/686 and 815/816) resulting in S1, S2, and S’ subunits. After cleavage, the byproduct S1 and membrane-anchored S2 binds to the angiotensin converting enzyme 2 (ACE2) receptor to enter into the cell (Figure 6). Our analysis identified more than 20 mutation sites in the S protein, and all of those mutations were nonsynonymous. Out of these 20 mutations, 13 were in the N-terminal domain (NTD) fragment, i.e., T95I, Q14H, S13I, T75I, H49Y, N211Y, D138H, V127F, P26, G75V, S255, Y248H, and S95F. The remaining other sites were located in different regions within the protein including L5F, D614G, G769V, E516Q, T791I, L518I etc (Figure 6). The fusion peptide region, S’ including heptad repeats HR1 and HR2 regions contains 3 mutated regions (G769V, T791I, A783S), 2(S939Y D936Y), 1(K1191N) respectively, The RdRp peptide region was highly conserved than others as only two mutation sites (E516Q, L518I) were found (Figure 6 and Supplementary Material S2).

**Figure 6.**
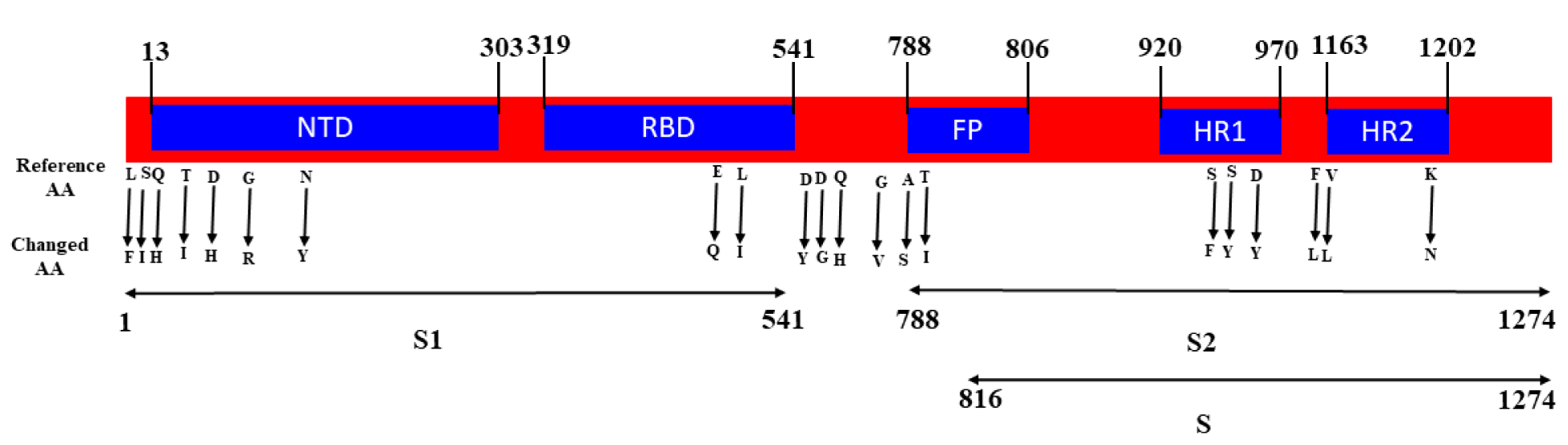
Mapping of mutations in different domains of spike protein. S1 and S2 are subdomains, N-terminal domain (NTD), C-terminal domain (CTD), Receptor binding domain (RBD), Fusion peptides(FP), Heptad repeats (HR1, HR2) regions, while S includes heptad repeats (HR1, HR2) regions. Changes in Amino acid (AA) sequence from the reference genome are shown by the arrow.

### 3.4 Sex and age based mutational accumulation analysis

The sex based unique mutation frequency during the entire period of analysis (13 weeks) was found to be the highest in man (70) than the women (36) (Table 2). Among the five segregated regions, R1 and R4 accumulated more mutations in man than the other regions. This is also true in case of females but with much lower frequency. Protein NSP10, Helicase, 3’ to 5’ exonuclease, ORF6, ORF7a had no mutation in case of females. NSP2, NSP3 RdRp, ORF8 and N proteins had unique mutations ranging between 2 and 6 amino acids per protein in the male originated virus (Table 1). Moreover, region 3 had the lowest unique mutation frequency in viral sequences retrieved from female patients. This aforementioned analysis data suggest differences in COVID-19 infection based on the sex of the infected individual. On the other hand, all age groups (4) accumulated the highest number of mutations in the virus genome R5, while the age group of 47-67 years harbors the highest number of mutational accumulation followed by the group of 26-46 years old (Figure 7). However, the age group 67 to 95 years old had the highest mutational frequency in R5, while a gradual increase of the mutational frequency was observed in case of age group 26-46 and 47 to 67 in all regions except R2. Most interestingly, age group 0 to 25 have approximately mutational accumulation rates similar to the age group 26 to 46 years old (Figure 7).

**Table 2:** Gender basic unique mutations of five regions occurred in more than one week in SARS-CoV-2 virus. Unique mutations were identified by removing redundant mutations that occurred in more than one week.

| Gender | Region 1 | Region 2 | Region 3 | Region 4 | Region 5 | Total |
| --- | --- | --- | --- | --- | --- | --- |

|  | Leader | NSP2 | NSP3 | NSP4 | NSP5 | NSP6 | NSP7 | NSP10 | RDRP | Helicase | 3' to 5'<br>exonuclease | EndoRNase | Spike | ORF3a | E | 6 | 7A | 8 | N |  |
| --- | --- | --- | --- | --- | --- | --- | --- | --- | --- | --- | --- | --- | --- | --- | --- | --- | --- | --- | --- | --- |
| Male | 1 | 6 | 15 | 2 | 4 | 4 | 0 | 1 | 3 | 3 | 2 | 2 | 8 | 9 | 1 | 1 | 2 | 4 | 2 | 70 |
|  | 22 |  |  | 10 |  |  |  | 9 |  |  |  | 19 |  |  | 10 |  |  |  |  |  |
| Female | 3 | 2 | 5 | 1 | 1 | 2 | 1 | 0 | 2 | 0 | 0 | 2 | 5 | 8 | 1 | 0 | 0 | 1 | 2 | 36 |
|  | 10 |  |  | 5 |  |  |  | 2 |  |  |  | 15 |  |  | 4 |  |  |  |  |  |

**Figure 7.**
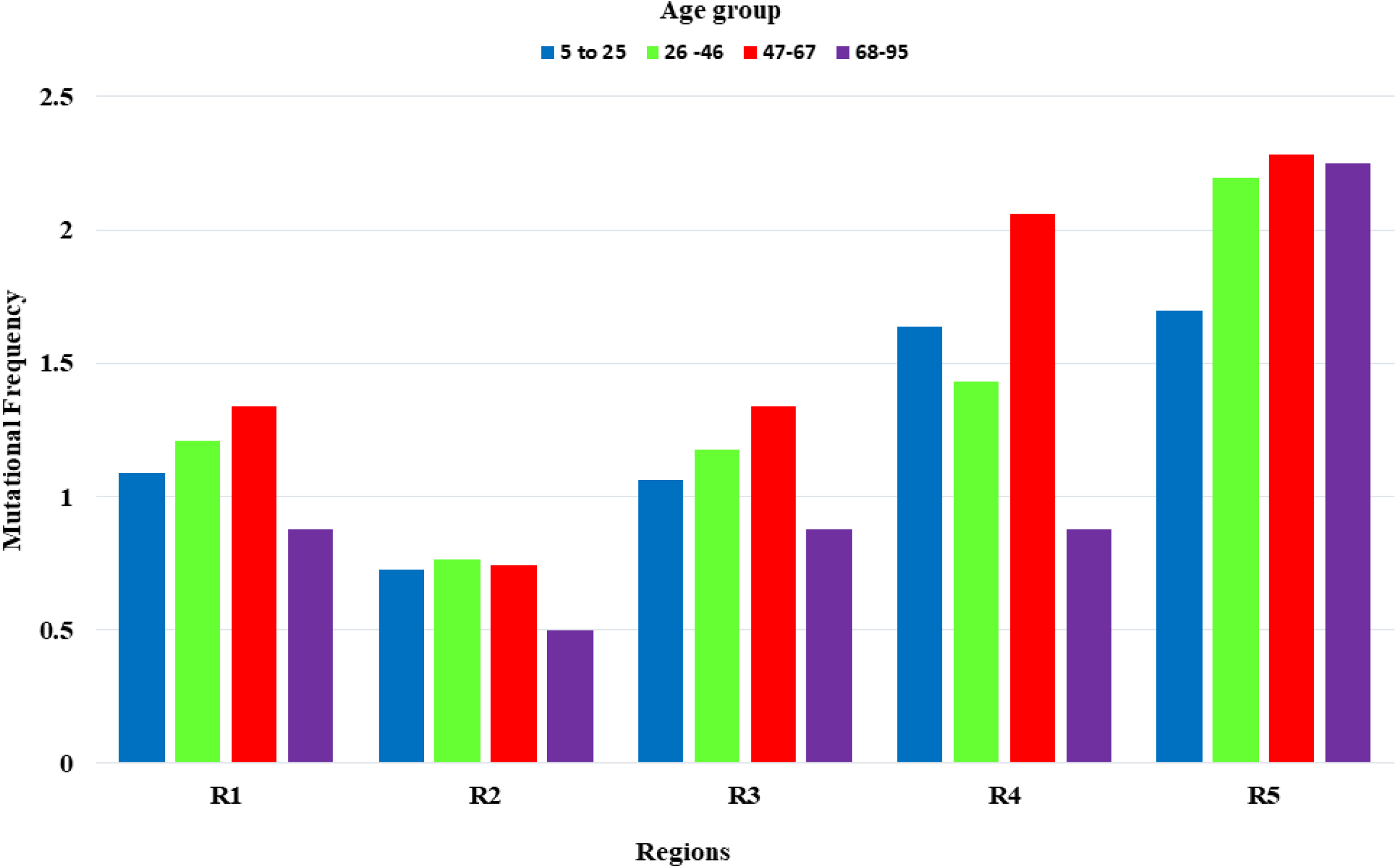
Age based mutational frequency of five genomic segments of SARS-CoV-2. Mutational frequency was calculated by the ratio of the number of total protein mutations and the number of genome sequences in each week. The SARS-CoV-2 genome was divided into five regions, which are represented as R1–R5. All the studied patients were segregated into 4 age groups (0 to 25 years-blue color, 26 to 46 years-green color, 47-67 years-red color, 68 to 95 years-violet color).

## 4. Discussion

Considering the lack of definitive drug and vaccine against COVID-19, studying SARS-CoV-2 genomes is of great importance to elucidate the molecular basis of pathogenesis and evolution for explaining differences in region specific mortality rates and individual dependent susceptibility to SARS-CoV-2. The current study of analyzing 198 high quality complete SARS-CoV-2 genomic sequences from Bangladesh revealed that the circulating strains are of many sub-lineages harboring the same ancestry as Wuhan virus. Although their direct evolution from that reference Wuhan virus was not found. So, it can be suggested that viruses from different regions other than China might also contribute to the SARS-CoV-2 outbreak in Bangladesh, which was also previously reported by Parvez et al. 2020; Hasan et al. 2020. Besides, that majority of the Bangladeshi isolates were found to fall within the clade B belonging to L type (Supplementary Table ST2). While these types were estimated to be more aggressive and capable of rapid transmission, human intervention had been reported to decrease the relative frequency of the L type (Tang et al., 2020). Similar type of isolates originating from type A subtype were also reported circulating into the European countries by Forster et al. in 2020, while a recent study reported the emergence of European and North American mutant variants in Southeast Asia including Bangladesh (Islam et al., 2020). However, mutational frequency analysis of the SARS-CoV-2 whole genomes has shown fluctuations of mutational frequency over time, which can be associated with the increase or decrease rate of infections among the population of Bangladesh (IEDCR, 2020). Among the 5 regions of SARS-CoV-2 genomes divided to determine the region of mutational hotspots, regions (R) 1, 4, 5 showed a greater tendency to accumulate mutations compared to region 2 and 3. On the other hand, the temporal profile of mutational analysis showed a higher mutational rate in weeks 7–10 and the mutational rate was increasing over time, which was also consistent with the study carried out in USA (Kaushal et al., 2020). Regions 2 and 3 of higher conservancy harbored NSP4, NSP 5, NSP6, NSP 7, NSP8, NSP 9, NSP10, and NSP11, while the conservative nature of these proteins was also reported previously (Kaushal et al., 2020; Korber et al., 2020). Besides, NSP9 and NSP11 had no record to accumulate any amino acid substitutions over a period of 11 weeks in USA, which was evident in our study in case of NSP9. However, only two amino acid substitutions were identified in NSP11 protein by Liang et al. in 2020. So, NSP9 and NSP11 could be potential targets for the treatment of SARS-CoV-2 to reduce local inflammation and tissue damage. Conversely, the amino acid mutation frequency was higher in NSP2, NSP3, RdRp, ORF3a, but the maximum number of mutations were detected in N (Nucleocapsid) and S (Spike) protein, which contradicts the finding of to the study carried out by Kaushal et al. in 2020 reporting higher mutation in the region of ORF8 and helicase. However, the mutational frequencies in these regions may positively facilitate the virus to adapt to not only external interactions with host cells, but also internal interactions within the host cells (Kim et al., 2020). While the mutation site of N protein does not elicit much antibody response, region 603-634 of the S protein of SARS has been shown to be a major immunodominant epitope in S protein (He et al., 2004). So, changes in this epitope by mutation could alter the sensitivity of the IgG/IgM tests conducted. These changes are actually due to the positive selection pressure in SARS-CoV-2 (Velazquez-Salinas et al., 2020, Benvenuto et al., 2020). Additionally ORF8, ORF7a and ORF7b showed mutations in many SARS-CoV-2 isolates, which might result in significant adaptation of coronavirus from human-to-human transmission as well as in contributing to the viral pathogenesis in the host by inhibiting bone marrow stromal antigen 2 (BST-2), which restricts the release of coronaviruses from affected cells (Decaro et al., 2020; Taylor et al., 2015). In this study, signature nonsynonymous mutations leading to amino acid changes of P323L in the RdRp was found (Supplementary Table ST2), which is involved in the replication of the viral genome. Moreover, D614G in the spike glycoprotein is also predominant in Bangladesh originated SARS-CoV-2 genome, which should be of urgent concern considering the dominance of this mutation globally since early February in Europe (Korber et al., 2020). Notably, the D614G mutation is close to the furin recognition site for cleavage of the spike protein, which plays an important role in virus entry. So, mutations in S protein including D614G need to be evaluated carefully, as S protein is essential for the entry of the virus in the host cell by binding to the ACE2 receptor leading to the escape from antibody inhibition allowing infected and recovered patients to become infected again (Huang et al., 2020) and these mutations may have resulted in the evolution of a new subtype with more transmissible ability (Maitra et al., 2020).

Several unique mutations in NSP3 followed by S, ORF3a, NSP2, RdRp, helicase, E and N protein were observed in this study, which may contribute to virulence, transmission, and pathogenicity during the epidemic (Consortium et al., 2004). Interestingly, accumulating mutations were found to be higher in males than female patients, which was also reported from other countries including Italy showing male to female ratio being 3:1 in Italy (Chakravarty et al., 2020). The mortality rate was also reported high in males compared to females from China showing 2.4 times higher mortality in males (Jian-Min et al., 2020), New York State of USA (42% females vs. 58% males (https://www.syracuse.com/coronavirus-ny/). However, age stratified mortality rate was also evident (Guilmoto et al., 2020), while the mutational accumulation in this study also showed an age-stratified pattern. The highest number of mutation accumulation was observed in age group of 47-67 years, followed by group of 26 to 46 years old which, might explain the infection rate in Bangladesh (IEDCR, 2020). On the other hand, the temporal analysis of mutation accumulation also showed that several mutations of SARS-CoV-2 proteins were persistent over the weeks of analysis, while some of the persistent mutations were reported previously from different countries, suggesting similarities in SARS-CoV-2 evolution across the world (Chan et al., 2020; Ou et al., 2020; Saha et al., 2020). Moreover, RdRp, E, and N genes are the target genes for designing primers and probes in RT-PCR-based SARS-CoV-2 diagnosis (Khailany et al., 2020), owing to their high sequence conservation. Although, it has not been known how the primer-template mismatches affect the accuracy and precision of the genetic diagnosis of COVID-19. So any kind of changes in this reasons might play a vital role in the diagnosis of the SARS-CoV-2 (Khailany et al., 2020). Unfortunately in our studied genome we have found majority of the genomic variation in the RdRp and N region which might be more alarming. In a study by Khan et al. in 2020 reported 7 set of genomic diagnostic assay out of 27 assay contain mismatches or mutation which is also supported by our studied results where 5 set of genomic diagnostic assay contain the following mismatches (Table 3) with having 100% in some cases which is very much alarming for the diagnostic procedure in Bangladesh.

**Table 3:**
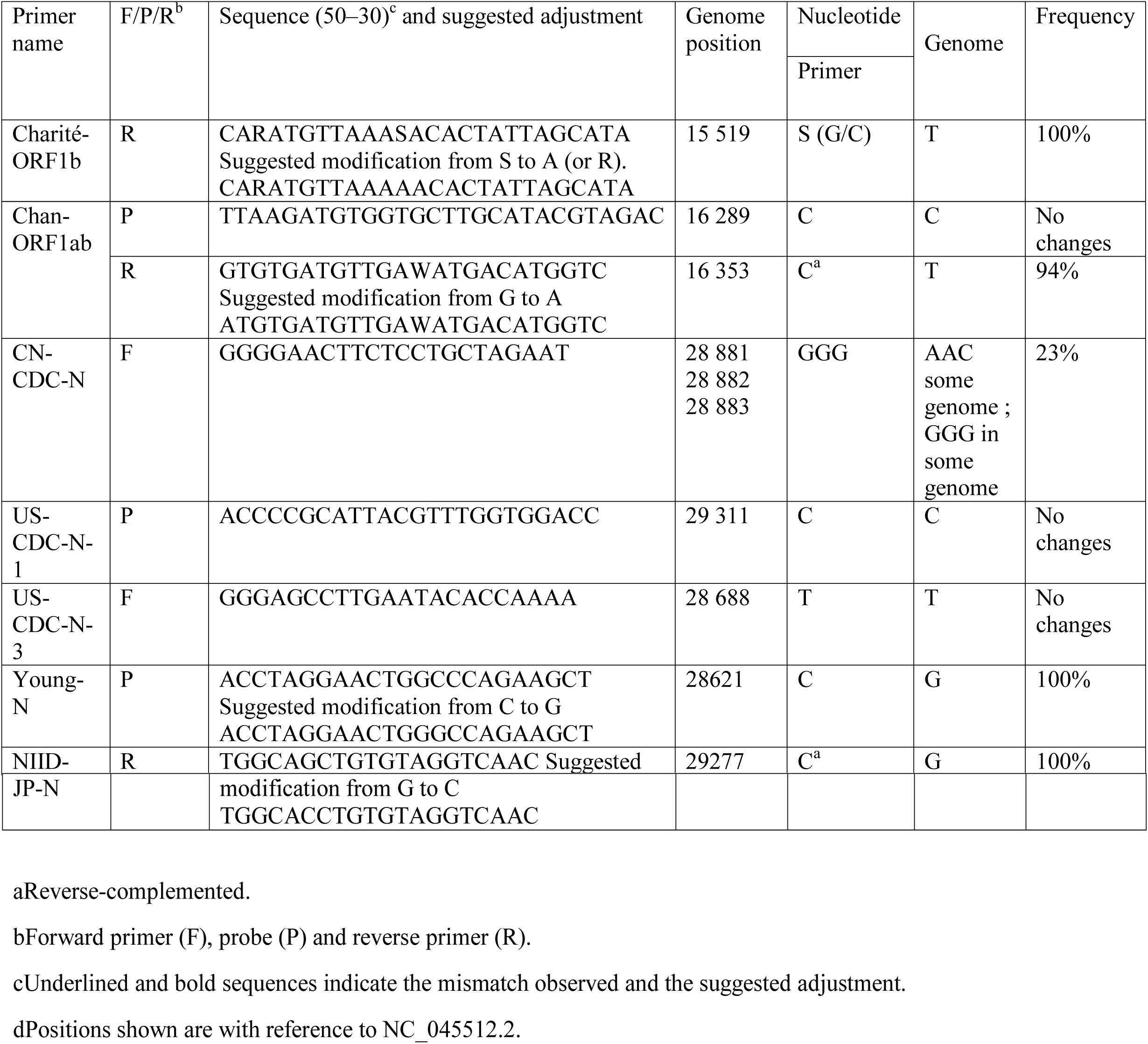
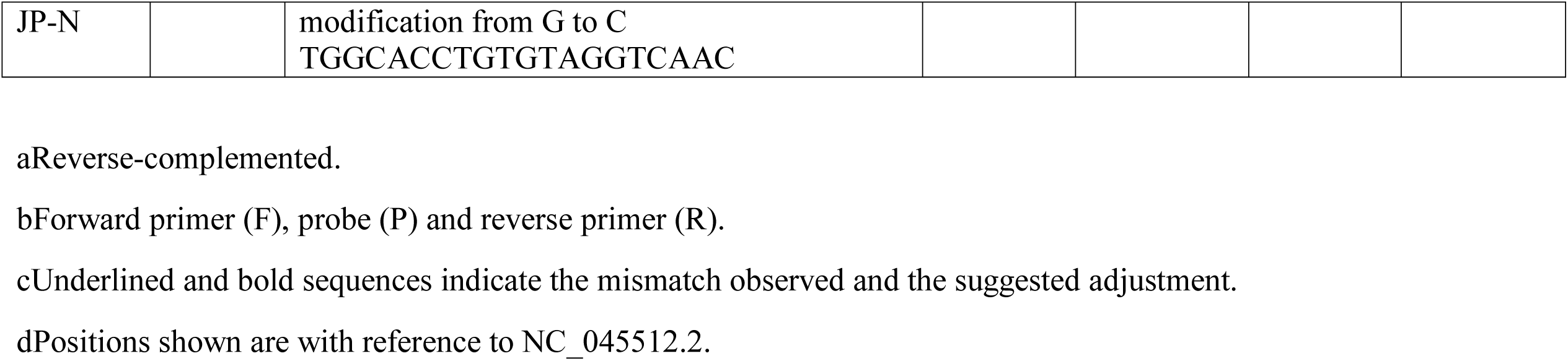
Summary of primer/probe mismatches with SARS-CoV-2 genome.

However, it appears that country and continent specific mutations are being accumulated in the SARS-CoV-2 genomes, while most of the vaccine attempts and diagnostic kits are based on the genome sequence of the original viral isolate from Wuhan. So, the region-specific mutations in SARS-CoV-2 genome may make these vaccines ineffective. Therefore, continuous monitoring of mutation accumulation and the consequences of these mutations on receptor binding affinity, genome replication and propagation ability, pathogenicity as well as the host-pathogen interaction need to be evaluated. On the other hand, RdRp, E, and N genes should be considered as the target genes for designing primers and probes in RT-PCR-based SARS-CoV-2 diagnosis, owing to their high sequence conservation.

## Author’s contributions

S., R.N.S., and I.I. carried out the studies (Data collection and data analysis). O.S. drafted the manuscript. O.S. and M.M.R. developed the hypothesis, supervised the whole work and M.M.R., M.S.H., I.I., N.N.R. and R.N.S. critically reviewed the drafted manuscript. All authors read and approved the final manuscript.

## Funding source

No Funding

### Acknowledgments

The authors would like to acknowledge Bangabandhu Science & Technology Fellowship Trust for supporting Otun Saha with PhD fellowship.

## Ethical Statement

The authors confirm that the ethical policy of the journal, as mention on the journal authors guideline page, have been adhere to no ethical approval was required as this did not collect any sample or questionnaires from animals and humans.

## Data Availability Statement

Complete genome sequences or SARS-CoV-2 are available in GISAID dataset along with references sequence was in the NCBI dataset. The accession number and sequences of the studied genomes are available in the Supplementary table 1 and Supplementary file 1.

## Abbreviations

COVID-19: Coronavirus Disease 2019
ACE2: angiotensin converting Enzyme 2
MSA: Multiple Sequence Alignment
SARS: Severe Acute Respiratory Syndrome
SARS-CoV-2: Severe Acute Respiratory Syndrome Coronavirus 2
SNP: Single Nucleotide Polymorphisms
UM: Unique Mutations
WHO: The World Health Organization
GISAID: Global Initiative on Sharing All Influenza Data.

## References

Andersen, K. G., Rambaut, A., Lipkin, W. I., Holmes, E. C., and Garry, R. F. (2020). The proximal origin of SARS-CoV-2. Nat. Med, 26, 450–452. doi: 10.1038/s41591-020-0820-9

Benvenuto, D.; Giovanetti, M.; Ciccozzi, A.; Spoto, S.; Angeletti, S.; Ciccozzi, M. (2020). The 2019-new coronavirus epidemic: Evidence for virus evolution. J. Med. Virol, 92, 455–459.

Ceraolo C, Giorgi FM. (2020). Genomic variance of the 2019-nCoV coronavirus. J Med Virol, 92(5), 522–8.

Chakravarty, D., Nair, S. S., Hammouda, N., Ratnani, P., Gharib, Y., Wagaskar, V., … & Tewari, A. K. (2020). Sex differences in SARS-CoV-2 infection rates and the potential link to prostate cancer. Communications biology, 3(1), 1–12.

Chan, J.F.; Kok, K.H.; Zhu, Z.; Chu, H.; To, K.K.; Yuan, S.; Yuen, K.Y. (2020). Genomic characterization of the 2019 novel human-pathogenic coronavirus isolated from a patient with atypical pneumonia after visiting Wuhan. Emerg. Microbes Infect, 9, 221–236.

Consortium, C.S.M.E. (2004). Molecular evolution of the SARS coronavirus during the course of the SARS epidemic in China. Science, 303, 1666–1669.

Coppee, F.; Lechien, J.R.; Decleves, A.E.; Tafforeau, L.; Saussez, S. (2020). Severe acute respiratory syndrome coronavirus 2: Virus mutations in specific European populations. New Microbes New Infect, 36, 100696.

da Silva, S. J. R., da Silva, C. T. A., Mendes, R. P. G., & Pena, L. (2020). Role of Nonstructural Proteins in the Pathogenesis of SARSLCoVL2. Journal of Medical Virology, 1–3.

Dawood, A. A. (2020). Mutated COVID-19, may foretells mankind in a great risk in the future. New Microbes and New Infections, 100673.

Decaro, N., & Lorusso, A. (2020). Novel human coronavirus (SARS-CoV-2): A lesson from animal coronaviruses. Veterinary Microbiology, 108693.

Fani M, Teimoori A, Ghafari S. (2020). Comparison of the COVID-2019 (SARS-CoV-2) pathogenesis with SARS-CoV and MERS-CoV infections. Future Virol, 10.2217/fvl-2020-0050. doi: 10.2217/fvl-2020-0050.

Fauver, J. R., Petrone, M. E., Hodcroft, E. B., Shioda, K., Ehrlich, H. Y., Watts, A. G., … & Razeq, J. (2020). Coast-to-coast spread of SARS-CoV-2 during the early epidemic in the United States. Cell, 181(5), 990–96.

Forster P, Forster L, Renfrew C, Forster M. (2020). Phylogenetic network analysis of SARS-CoV-2 genomes. Proc Natl Acad Sci, 117(17), 9241. doi: 10.1073/pnas.2004999117

Guilmoto, C. Z. (2020). COVID-19 death rates by age and sex and the resulting mortality vulnerability of countries and regions in the world. medRxiv.

Hasan, S., Khan, S., Ahsan, G. U., & Hossain, M. M. (2020). Genome Analysis of SARS-CoV-2 Isolate from Bangladesh. BioRxiv.

He Y, Zhou Y, Wu H, Luo B, Chen J, et al. (2004). Identification of immunodominant sites on the Spike protein of severe acute respiratory syndrome (SARS) coronavirus: Implication for developing SARS diagnostics and vaccines. J. Immunol, 173, 4050–4057. doi: 10.4049/jimmunol.173.6.4050.

Hon, C. C., Lam, T. Y., Shi, Z. L., Drummond, A. J., Yip, C. W., Zeng, F., … & Leung, F. C. C. (2008). Evidence of the recombinant origin of a bat severe acute respiratory syndrome (SARS)-like coronavirus and its implications on the direct ancestor of SARS coronavirus. Journal of virology, 82(4), 1819–1826.

Huang, A. T., Garcia-Carreras, B., Hitchings, M. D., Yang, B., Katzelnick, L. C., Rattigan, S. M., … & Lessler, J. (2020). A systematic review of antibody mediated immunity to coronaviruses: antibody kinetics, correlates of protection, and association of antibody responses with severity of disease. medRxiv.

Institute of Epidemiology, Disease Control and Research. COVID-19 status Bangladesh. [Cited 2020 August 14 2020]. Available from: https://www.iedcr.gov.bd/

Islam, M. M., Rakhi, N. N., Islam, O. K., Saha, O., & Rahaman, M. M. (2020). Challenges to be considered to evaluate the COVID-19 preparedness and outcome in Bangladesh. International Journal of Healthcare Management, 1–2.

Jian-Min Jin, P. B. et al. (2020). Gender differences in patients with COVID-19: focus on severity and mortality. Front. Public Health, 8, 152. https://doi.org/10.3389/fpubh.2020.00152 (2020).

Kaushal, N., Gupta, Y., Goyal, M., Khaiboullina, S. F., Baranwal, M., & Verma, S. C. (2020). Mutational Frequencies of SARS-CoV-2 Genome during the Beginning Months of the Outbreak in USA. Pathogens, 9(7), 565.

Khailany, R. A., Safdar, M., & Ozaslan, M. (2020). Genomic characterization of a novel SARS-CoV-2. Gene reports, 1006, 82

Khan, K. A., & Cheung, P (2020). Presence of mismatches between diagnostic PCR assays and coronavirus SARS-CoV-2 genome. Royal Society Open Science. 7(6), 200636.

Kim, J. S., Jang, J. H., Kim, J. M., Chung, Y. S., Yoo, C. K., & Han, M. G. (2020). Genome-Wide Identification and Characterization of Point Mutations in the SARS-CoV-2 Genome. Osong Public Health and Research Perspectives, 11(3), 101.

Korber, B., Fischer, W., Gnanakaran, S. G., Yoon, H., Theiler, J., Abfalterer, W., … & Partridge, D. G. (2020). Spike mutation pipeline reveals the emergence of a more transmissible form of SARS-CoV-2. bioRxiv.

Kumar, S., Stecher, G., & Tamura, K. (2016). MEGA7: molecular evolutionary genetics analysis version 7.0 for bigger datasets. Mol. Biol. Evol, 33(7), 1870–1874

Liang, Q.; Li, J.; Guo, M.; Tian, X.; Liu, C.; Wang, X.; Yang, X.; Wu, P.; Xiao, Z.; Qu, Y. (2020). Virus-host interactome and proteomic survey of PMBCs from COVID-19 patients reveal potential virulence factors influencing SARS-CoV-2 pathogenesis. bioRxiv.

Lv, L., Li, G., Chen, J., Liang, X., and Li, Y. (2020). Comparative genomic analysis revealed specific mutation pattern between human coronavirus SARS-CoV-2 and Bat-SARSr-CoV RaTG13. BioRxiv

Maitra A, Sarkar MC, Raheja H, et al. (2020). Mutations in SARS-CoV-2 viral RNA identified in Eastern India: Possible implications for the ongoing outbreak in India and impact on viral structure and host susceptibility. J Biosci, 45(1):76. doi: 10.1007/s12038-020-00046-1

Mim, M. A., Rakhi, N. N., Saha, O., & Rahaman, M. M. (2020). Recommendation of fecal specimen for routine molecular detection of SARS-CoV-2 and for COVID-19 discharge criteria. Pathogens and global health, 1–2.

Ou, J.; Zhou, Z.; Dai, R.; Zhang, J.; Lan, W.; Zhao, S.; Wu, J.; Seto, D.; Cui, L.; Zhang, G. (2020). Emergence of RBD mutations in circulating SARS-CoV-2 strains enhancing the structural stability and human ACE2 receptor affinity of the spike protein. BioRxiv.

Parvez, M. S. A., Rahman, M. M., Morshed, M. N., Rahman, D., Anwar, S., & Hosen, M. J. (2020). Genetic analysis of SARS-CoV-2 isolates collected from Bangladesh: insights into the origin, mutation spectrum, and possible pathomechanism. bioRxiv.

Rahaman, M. M., Saha, O., Rakhi, N. N., Chowdhury, M. M. K., Sammonds, P., & Kamal, A. M. (2020). Overlapping of locust swarms with COVID-19 pandemic: a cascading disaster for Africa. Pathogens and Global Health, 1–2.

Rahman, M. S., Hoque, M. N., Islam, M. R., Akter, S., Rubayet-Ul-Alam, A. S. M., Siddique, M. A. & Hossain, M. A. (2020). Epitope-based chimeric peptide vaccine design against S, M and E proteins of SARS-CoV-2 etiologic agent of global pandemic COVID-19: an in silico approach. PeerJ, 8, e9572.

Saha, O., Rakhi, N. N., Towhid, S. T., & Rahaman, M. M. (2020). Reactivation of Severe Acute Respiratory Coronavirus-2 (SARS-CoV-2): Hoax or hurdle?. International Journal of Healthcare Management, 1–2.

Saha, P.; Banerjee, A.K.; Tripathi, P.P.; Srivastava, A.K.; Ray, U. (2020). A virus that has gone viral: Amino acid mutation in S protein of Indian isolate of Coronavirus COVID-19 might impact receptor binding, and thus, infectivity. Biosci. Rep, 40.

Saitou, N., & Nei, M. (1987). The neighbor-joining method: a new method for reconstructing phylogenetic trees. Mol. Biol. Evol, 4(4), 406–425.

Sevajol, M., Subissi, L., Decroly, E., Canard, B., & Imbert, I. (2014). Insights into RNA synthesis, capping, and proofreading mechanisms of SARS-coronavirus. Virus research, 194, 90–99.

Shen, Z., Xiao, Y., Kang, L., Ma, W., Shi, L., Zhang, L., et al. (2020). Genomic diversity of SARS-CoV-2 in coronavirus disease 2019 patients. Clin. Infect. Dis. ciaa203. doi: 10.1093/cid/ciaa203.

Su, Y. C., Anderson, D. E., Young, B. E., Linster, M., Zhu, F., Jayakumar, J., & Chia, W. N. (2020). Discovery and Genomic Characterization of a 382-Nucleotide Deletion in ORF7b and ORF8 during the Early Evolution of SARS-CoV-2. Mbio, 11(4).

Tai, W., He, L., Zhang, X., Pu, J., Voronin, D., Jiang, S., … & Du, L. (2020). Characterization of the receptor-binding domain (RBD) of 2019 novel coronavirus: implication for development of RBD protein as a viral attachment inhibitor and vaccine. Cellular & molecular immunology, 17(6), 613–620.

Tang, X. C., Agnihothram, S. S., Jiao, Y., Stanhope, J., Graham, R. L., Peterson, E. C., … & Baric, R. S. (2014). Identification of human neutralizing antibodies against MERS-CoV and their role in virus adaptive evolution. Proceedings of the National Academy of Sciences, 111(19), E2018–E2026.

Tang X, Wu C, Li X, et al. (2020). On the origin and continuing evolution of SARS-CoV-2. Natl Sci Rev. (nwaa036). doi: 10.1093/nsr/nwaa036.

Taylor, J. K., Coleman, C. M., Postel, S., Sisk, J. M., Bernbaum, J. G., Venkataraman, T., … & 290 Frieman, M. B. (2015). Severe acute respiratory syndrome coronavirus ORF7a inhibits bone 291 marrow stromal antigen 2 virion tethering through a novel mechanism of glycosylation 292 interference. Journal of virology, 89(23), 11820–11833.

Velazquez-Salinas, L.; Zarate, S.; Eberl, S.; Gladue, D.P.; Novella, I.; Borca, M.V. (2020). Positive selection of ORF3a and ORF8 genes drives the evolution of SARS-CoV-2 during the 2020 COVID-19 pandemic. BioRxiv.

Waterhouse, A.M.; Procter, J.B.; Martin, D.M.; Clamp, M.; Barton, G.J. (2009). Jalview Version 2—A multiple sequence alignment editor and analysis workbench. Bioinformatics, 25, 1189–1191.

Wu, C., Liu, Y., Yang, Y., Zhang, P., Zhong, W., Wang, Y., et al. (2020). Analysis of therapeutic targets for SARS-CoV-2 and discovery of potential drugs by computational methods. Acta Pharm. Sin. B 10, 766–788. doi: 10.1016/j.apsb.2020.02.008

Wu, F., Zhao, S., Yu, B., Chen, Y.M., Wang, W., Song, Z.G., Hu, Y., Tao, Z.W., Tian, J.H., Pei, Y.Y., et al. (2020). A new coronavirus associated with human respiratory disease in China. Nature 579, 265–269.

Yu, W.-B. (2020). Decoding evolution and transmissions of novel pneumonia coronavirus (SARS-CoV-2) using the whole genomic data Comparative analyses of the chloroplast genome in carnivorous plants View project. ChinaXriv.

Zhao, S., and Chen, H. (2020). Modeling the epidemic dynamics and control of COVID-19 outbreak in China. Quant. Biol, 8, 11–19. doi: 10.1007/s40484-020-0199-0

Zhou, Y., Hou, Y., Shen, J., Huang, Y., Martin, W., and Cheng, F. (2020). Network-based drug repurposing for novel coronavirus 2019-nCoV/SARS-CoV-2. Cell Discov. 6:14. doi: 10.1038/s41421-020-0153-3.

